# Winter activity of boreal bats

**DOI:** 10.1101/2020.10.19.345124

**Authors:** Anna S. Blomberg, Ville Vasko, Melissa B. Meierhofer, Joseph S. Johnson, Tapio Eeva, Thomas M. Lilley

## Abstract

Natural hibernation sites used by bats in areas that lack cave features have long remained unresolved. To investigate hibernation site selection and winter activity of boreal bats, we recorded bat calls using passive acoustic monitoring on 16 sites. These sites included four rock outcrops with crevices and cave features, three glacial erratics or boulder fields, three ancient shores, three root cellars and three control sites where we did not expect bats to be overwintering. Our results revealed echolocation calls of *Eptesicus nilssonii*, *Plecotus auritus* and *Myotis* sp. We recorded significantly more activity near rock outcrops and root cellars compared to other habitats. We also found that ambient temperature had a positive effect on bat activity and found evidence that *P. auritus* may be using low barometric pressure as a proxy for suitable foraging conditions during the winter. Our results suggest that rock outcrops may be more important to bats than previously acknowledged, highlighting the need to take these sites in account in planning of conservation measures. Furthermore, our findings underline the suitability of using acoustic monitoring in homing on hibernation sites that are not otherwise accessible.

## INTRODUCTION

Insectivorous bats living at high latitudes face enormous fluctuations in the seasonal availability of food. Bats can respond to this challenge either by migrating to warmer areas, where food is occasionally available throughout the winter and hibernation is less risky (Popa-Lisseanu and Voigt 2009), or they can hibernate *in situ*, by utilizing fat reserves accumulated before the winter (Geiser 2013). The hibernation period of temperate bats can last more than eight months (Norquay and Willis 2014) and is elapsed in underground hibernation sites, hibernacula. Hibernation consists of extended bouts of torpor, during which bats lower their body temperature close to ambient temperature of the hibernaculum and decrease their metabolic rate (Geiser 2004; Guppy and Withers 2007). Torpor is interrupted with euthermic bouts — arousals — which allow the bat to counter the ecological and physiological costs caused by torpor (Thomas et al. 1990; Humphries et al. 2003; Boyles and Brack 2009; Bouma et al. 2010; Lilley et al. 2017). Arousals are energetically costly, and amount to 80–90% of total energy expenditure during the winter (Thomas et al. 1990). Due to the long winters at high latitudes, bats must shift their microclimate preference towards colder ambient temperatures and longer torpor bouts to conserve energy (Dunbar and Brigham 2010). Therefore, results of winter activity studies conducted in milder climates, where food is available throughout the winter, are not applicable in the north (Hope and Jones 2012).

During arousals, bats also have an opportunity to drink, copulate and switch places within the hibernaculum. Sometimes bats also leave the hibernation site entirely. While information on the activities that bats undertake outside the hibernaculum is scarce (Boyles et al. 2006), foraging has often been suggested as the primary driver for winter activity (Avery 1985; Dunbar et al. 2007; Zahn and Kriner 2016). Several studies have reported a positive relationship between bat activity and temperature (Avery 1985; Zahn and Kriner 2016; Klüg-Baerwald et al. 2016), as warm ambient temperatures may enable foraging while also minimizing the energetic costs of activity. However, it is unlikely that bats occurring at northern latitudes are able to feed during the coldest months, as there are almost no nocturnal aerial insects available. Bats may also need to relocate to a new hibernaculum during hibernation, which can be due to disturbance, unfavourable changes in abiotic conditions, or a shift in balance between the need to save energy versus minimize cost of torpor (Boyles et al. 2006; Masing and Lutsar 2007; Wermundsen and Siivonen 2010; Johnson et al. 2016). Masing and Lutsar (2007) reported that ambient winter temperatures that are low enough to reduce the temperature inside the hibernacula, caused *Eptesicus nilssonii* to arouse and switch to another hibernaculum, while *Myotis daubentonii*, *M. nattereri* and *Plecotus auritus* found in similar open hibernacula often froze to death.

If activity outside the hibernacula is due to foraging, bats should time their activity with elevated outside hibernacula ambient temperatures. However, most of the time hibernacula are well protected against changes in outside temperatures and therefore, bats cannot sense if it were to fluctuate. Unlike the outside temperature, changes in barometric pressure can be detected from inside the hibernaculum. Bats, like many other mammals, can sense changes in barometric pressure and they are known to utilize these cues in many circumstances (Paige 1995; Cryan and Brown 2007; Czenze and Willis 2015). Studies on the effect of barometric pressure on bat activity have reported conflicting results, with some studies describing increased activity with falling or low barometric pressure (Paige 1995; Czenze and Willis 2015), and other studies reporting a positive correlation between bat activity and barometric pressure (Berková and Zukal 2010; Bender and Hartman 2015). The different outcomes likely stem from methodological differences, as well as geographical and seasonal factors affecting both bat ecology and the relationship between weather conditions and barometric pressure.

Due to the geology of the Svecofennican orogeny, the majority of all previously known hibernacula in Fennoscandia are man-made (e.g., bunkers, cellars and mines) while very little is known about the use of natural formations as hibernacula. A telemetry study carried out in Norway found *Eptesicus nilssonii* and *Myotis mystacinus* hibernating in rock scree and rock crevices even though there were also man-made structures suitable for hibernation available (Michaelsen et al. 2013). Studies in North America have found bats using rock crevices for hibernation (Neubaum et al. 2006; Perry et al. 2010; Johnson et al. 2016; Lemen et al. 2017; Klüg-Baerwald et al. 2017; Neubaum 2018). Johnson et al. (2016) documented bats using ground level rock crevices, rock walls, scree slopes and rock fields also during late autumn. Currently there are no estimates on the number of bats in Finland but, given that only some thousands of bats are observed during the census each winter, it is likely that most bats hibernate in yet unidentified sites. Due to the glacial history and geology of Finland, there is a strong likelihood these hibernation sites are associated with a variety of rock formations. Because of the probability of attempting to forage or shift hibernacula during the hibernation period described above, acoustic monitoring could provide a means to gather information on the nature of these sites.

Of the eight bat species known to overwinter in Finland (Siivonen and Wermundsen 2008; Wermundsen and Siivonen 2010; Blomberg et al. 2020), five, including *E. nilssonii*, *P. auritus*, *Myotis daubentonii*, *M. brandtii* and *M. mystacinus* are considered common in South-Western Finland, where we conducted our study (Tidenberg et al. 2019; Vasko et al. 2020). In order to study the use of natural rock formations as hibernacula and factors affecting the winter activity of these boreal bats, we conducted acoustic monitoring of bat activity at locations we suspected were suitable hibernation sites. Our monitoring was conducted across all seasons to highlight winter activity at the sites. We selected our sites based on the findings in Norway (Michaelsen et al. 2013) and North America (Lausen and Barclay 2006; Johnson et al. 2016; Klüg-Baerwald et al. 2017), as well as anecdotal evidence on bats hibernating in boulder fields in Ostrobothnia and our previous knowledge of root cellars and some caves being used by bats in the winter. The abiotic conditions of the hibernacula affect the overwintering success of bats (Boyles and Brack 2009). Finding a suitable hibernaculum is vital for the survival of bats in colder climates, which is why we can predict that the location of hibernaculum determines much of the spatial distribution of bats during the winter. This is most likely the case especially in the north, where bats are unlikely to forage during the winter. Therefore, we consider bat activity at a given location to be a proxy for hibernation sites.

We predict that we would detect more bat activity during the winter months at our selected, potential hibernation sites compared to sites that do not have geological formations or structures suitable for hibernation. We also predict that bats will be more active during warm nights, and that low barometric pressure increases bat activity, as during the winter it can function as a proxy for mild outside temperature, which bats can sense from within their hibernacula (Paige 1995; Czenze and Willis 2015). Further, we predict *E. nilssonii* and *P. auritus* to be more active during the winter compared to *Myotis* species, as both are larger in size allowing them to allocate more fat resources for activity.

## MATERIALS AND METHODS

### Acoustic monitoring

We monitored year-round bat activity at 16 sites in southwestern Finland using AnaBat SD2 (Titley Scientific) and SongMeter SM2+ BAT (Wildlife Acoustics) passive detectors from 12 November 2017 to 31 April 2019. Monitoring sites selected included three ancient shores, four rock outcrops with caves and/or crack features (Fig 1), three glacial erratic formations or boulder fields, and three root cellars constructed by humans, of which the latter were known to be used by bats during the winter as hibernation sites. Ancient shores are post-glacial rebounds, upthrusts, where the continuously rising land causes the sea to retreat further from the continent leaving rocks and pebbles where the seashore was historically. The group consisting of glacial erratics or boulder fields included two glacial erratics, i.e. glacially-deposited rocks differing the size and type of rock native to the area in which it rests, and one man-made yet parallel boulder field. Three feeding sites used by bats in the summer were used as control sites for the study (Fig 2). We selected all sites based on their potential suitability for bats and accessibility for research purposes, because during the coldest and darkest period of winter, the batteries in the devices had to be changed every other week. At the sites, we placed microphones at 1.5 m height on a tree or a pole in semi-open environment with enough flying space for the bats.

**Fig 1.**
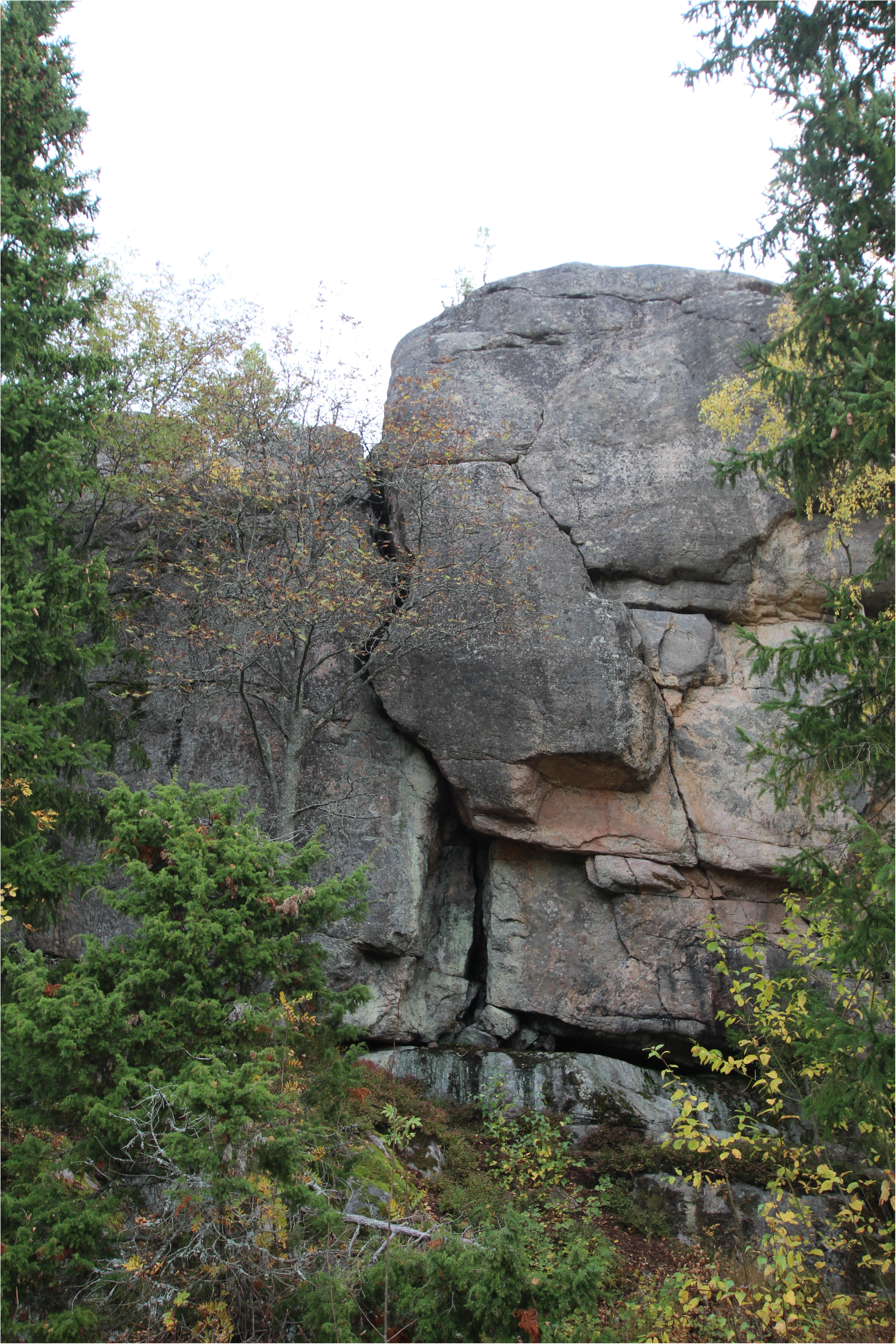
Acoustic monitoring during the winter months revealed a significant amount of bat activity in the vicinity of rock outcrops, suggesting that they are important hibernation sites for bats.

**Fig 2.**
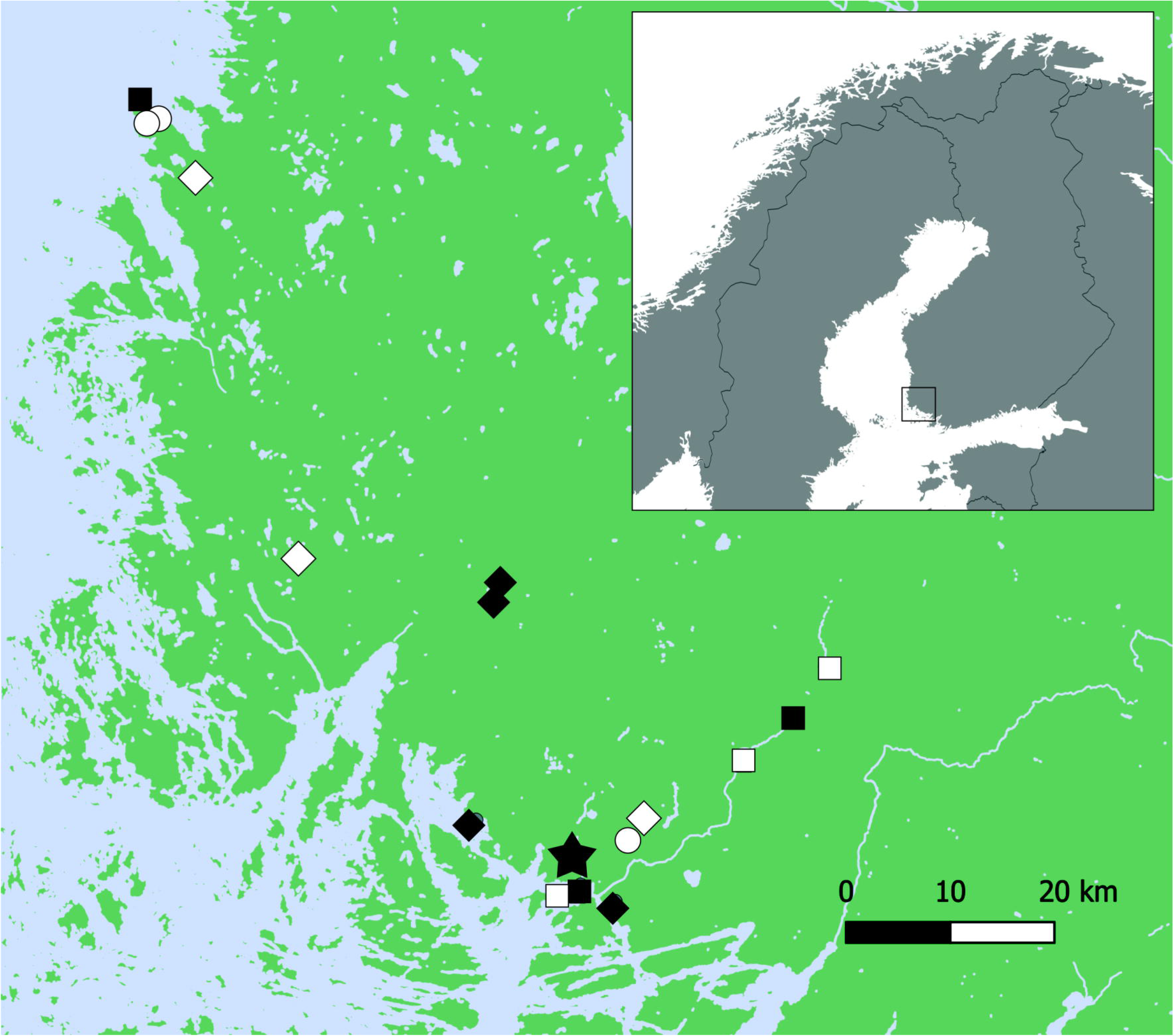
Map of study area. White squares = control sites, black squares = root cellars, white circles = ancient shores, black diamonds = outcrops, white diamonds = glacial erratics, star = weather station.

We programmed the SongMeter-passive detectors to begin monitoring half an hour before sunset and end monitoring half an hour after the sunrise. We adjusted the monitoring schedule of the AnaBat-passive detectors manually each month so that it matched the schedule of the SongMeters. We collected the data in WAV-format (SongMeters) and ZeroCrossing-format (AnaBats). Due to the harsh weather conditions, all multidirectional ultrasound microphones were fitted with a protective covering to keep them from getting wet or covered with snow. We used SMX-US microphones for the SongMeters and tested their sensitivity both before and after the monitoring. Microphones with sensitivity below –15dB at 40 kHz were not used, as recommended by the manufacturer. The AnaBats were tested by comparing them with a SongMeter in a side-by-side setup for two nights before and after the study. We obtained weather data from the Finnish Meteorological Institute. For the analysis, we used weather data from the closest weather station to the survey sites (Fig 2).

### Species identification

We used Kaleidoscope Pro (Version 5.3.8., Wildlife Acoustics) for filtering and organizing the data. We chose not to discard non-bat (noise) files suggested by the program from wintertime to ensure that all recorded bat calls were included in the analysis. From the summer months, noise files were discarded, because checking them manually would have been too laborious. After initial filtering, we then manually viewed acoustic files and determined species. Due to the uncertainty of identifying *Myotis* to a species level, we chose to pool all recordings of the genus *Myotis* for analysis.

### Statistical analysis

We considered winter months as those from November–March (Finnish Meteorological Institute). All together, we recorded 53 656 hours during the two winters for all sites. We gathered 35 minutes of *Myotis* activity, 182 minutes of *E. nilssonii* activity and 66 minutes of *P. auritus* activity. We investigated the differences in bat activity between sites, habitats and species during the winter with a generalized linear mixed model (Proc GLIMMIX in SAS, SAS Institute Inc. 2013). In the model, we used normal error distribution with identity link function. For each site, we calculated a monthly activity index for *E. nilssonii, P. auritus* and pooled *Myotis* species by dividing the number of active minutes (minutes with bat activity) by the number of total recorded minutes. We used this activity index with a log10-transformation as the response variable and species, habitat type and site nested under habitat type as fixed effects. In cases where we recorded zero minutes during a month at a site, we replaced the zero with 0.001 minutes to conduct the log10-transformation for the activity index. We also investigated the differences in activity between habitats and species using the Tukey-Kramer method.

Furthermore, we investigated the effect of ambient temperature and barometric pressure on the activity of *E. nilssonii*, *P. auritus* and the pooled set of *Myotis* -species, and all bat activity (all species pooled) using generalized linear models. We used the mean ambient temperature and mean barometric pressure of the previous 24 hours, as well as their interaction as explanatory variables. We used mean barometric pressure of the previous 24 hours, as low barometric pressure in the winter brings cloudy and rainy, yet mild weather to our study area. We standardized all fixed factors prior to analysis and checked for multicollinearity using variance inflation factors (VIFs). VIFs were < 3 in all instances, suggesting that no or very minimal multicollinearity between fixed factors existed. We used backward stepwise model selection, retaining the variables that produced the lowest AIC-values, to select the best model for each species and species group. We considered *P*-values < 0.05 significant for all tests.

## RESULTS

Levels of bat activity differed between habitats (F_4, 405_=9.91; *P* < 0.001), species (F_2, 405_ =19.29; *P*<0.001) and sites (F_11, 405_=2.11; *P*=0.019). Most bat activity during the winter took place at rock outcrops (Table 1). Rock outcrops had significantly more bat activity than ancient shores (t=4.51; DF=405; p<0.001), glacial erratics (t=4.19; DF=405; p<0.001) and control sites (t=5.09; DF=405; p<0.001). Second highest activity was measured outside root cellars, in which we previously knew bats were overwintering. Root cellars had significantly more activity in the winter than control sites, which were selected because they were known foraging areas in the summer (t=3.29; DF=405; p=0.01). The difference in activity between root cellars and ancient shores approached statistical significance (t=2.67; DF=405; p=0.06). Furthermore, our results revealed significantly more calls of *E. nilssonii* than *P. auritus* (t=5.17; DF=405; p<0.001) and *Myotis* species (t=5.56; DF=405; p<0.001; Table 2).

**Table 1.** Differences of LS-means for total bat activity measured at different habitats during the winter. Bat activity was measured at each site with a monthly activity index (minutes with recordings/number of recorded minutes).

| Habitat | Habitat | Estimate | SE | DF | t Value | Pr > \|t\| | Adj P | Alpha | Lower | Upper | Adj<br>Lower | Adj<br>Upper |
| --- | --- | --- | --- | --- | --- | --- | --- | --- | --- | --- | --- | --- |
| Ancient shore | Outcrop | -0.991 | 0.219 | 405 | -4.51 | <0.001 | <0.001 | 0.05 | -1.423 | -0.559 | -1.593 | -0.389 |
| Ancient shore | Glacial erratic | -0.034 | 0.243 | 405 | -0.14 | 0.889 | 0.910 | 0.05 | -0.513 | 0.444 | -0.701 | 0.633 |
| Ancient shore | Root cellar | -0.738 | 0.276 | 405 | -2.67 | 0.008 | 0.060 | 0.05 | -1.280 | -0.195 | -1.494 | 0.020 |
| Ancient shore | Control site | 0.217 | 0.252 | 405 | 0.86 | 0.390 | 0.911 | 0.05 | -0.278 | 0.711 | -0.473 | 0.906 |
| Outcrop | Glacial erratic | 0.957 | 0.228 | 405 | 4.19 | <0.001 | <0.001 | 0.05 | 0.508 | 1.406 | 0.331 | 1.582 |
| Outcrop | Root cellar | 0.253 | 0.263 | 405 | 0.96 | 0.336 | 0.871 | 0.05 | -0.263 | 0.770 | -0.467 | 0.974 |
| Outcrop | Control site | 1.207 | 0.237 | 405 | 5.09 | <0.001 | <0.001 | 0.05 | 0.741 | 1.674 | 0.558 | 1.857 |
| Glacial erratic | Root cellar | -0.703 | 0.283 | 405 | -2.49 | 0.013 | 0.096 | 0.05 | -1.260 | -0.147 | -1.479 | 0.072 |
| Glacial erratic | Control site | 0.251 | 0.259 | 405 | 0.97 | 0.334 | 0.870 | 0.05 | -0.259 | 0.760 | -0.459 | 0.961 |
| Root cellar | Control site | 0.954 | 0.290 | 405 | 3.29 | 0.001 | 0.010 | 0.05 | 0.384 | 1.524 | 0.159 | 1.749 |

**Table 2.**
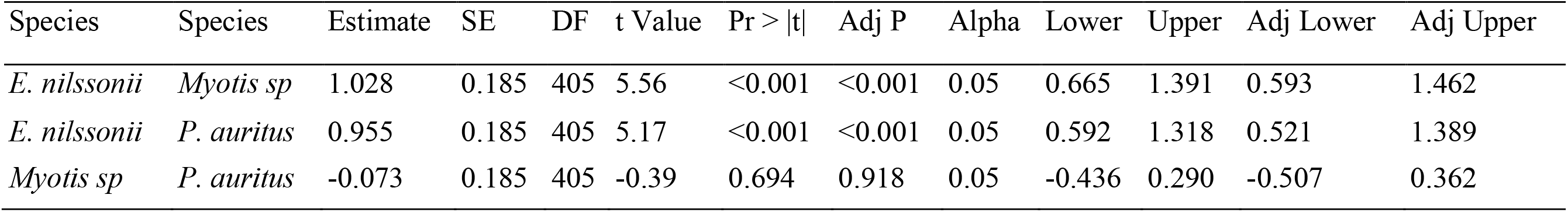
Differences of LS-means for monthly activity indexes of *Eptesicus nilssonii*, *Plecotus auritus* and a pooled set of *Myotis* species (*M. daubentonii*, *M. brandtii*, *M. mystacinus, M. nattereri*) recorded during the winter.

Total bat activity increased with higher ambient temperature and barometric pressure (Table 3). However, we found that total activity decreased faster with decreasing temperatures if barometric pressure was low (Fig. 3a). Activity of *E. nilssonii* was positively affected by increasing barometric pressure and ambient temperature (Table 3). Similar to *E. nilssonii*, the activity of *P. auritus* increased with higher mean ambient temperature (Table 3). The effect of temperature on the activity of *P. auritus* was stronger at low barometric pressure (Fig 3b). Contrary to *P. auritus*, the activity of *Myotis* species increased more steeply with ambient temperature at high barometric pressure (Fig 3c). The lowest hourly temperature where a bat recording was made, was −9,8 °C (*E. nilssonii*). The lowest temperature for the *Myotis* and *P. auritus* were −3,1 and −5,6 °C, respectively.

**Table 3.** Results of generalized linear model on the effect of weather conditions on total bat activity (n=318 minutes with activity), activity of *Eptesicus nilssonii* (N=182 minutes with activity), activity of *Plecotus auritus* (N=66 minutes with activity) and activity of *Myotis* sp. (pooled for four species; *M. daubentonii*, *M. brandtii*, *M. mystacinus*, *M. nattereri,* N=35 minutes with activity).

| Species | Weather variable | Estimate | Std. Error | z value | Pr(>\|z\|) |
| --- | --- | --- | --- | --- | --- |
| Total bat activity | (Intercept) | -2.092 | 0.078 | -26.650 | <0.001 |
| Total bat activity | Temperature | 0.982 | 0.092 | 10.649 | <0.001 |
| Total bat activity | Barometric pressure | 0.292 | 0.073 | 3.996 | <0.001 |
| Total bat activity | Temperature:Pressure | -0.166 | 0.074 | -2.240 | 0.025 |
| *E. nilssonii* | (Intercept) | -3.281 | 0.128 | -25.542 | <0.001 |
| *E. nilssonii* | Temperature | 0.859 | 0.139 | 6.192 | <0.001 |
| *E. nilssonii* | Barometric pressure | 0.232 | 0.123 | 1.880 | 0.06 |
| *P. auritus* | (Intercept) | -5.319 | 0.444 | -11.982 | <0.001 |
| *P. auritus* | Temperature | 2.010 | 0.529 | 3.800 | <0.001 |
| *P. auritus* | Barometric pressure | 0.910 | 0.359 | 2.536 | 0.01 |
| *P. auritus* | Temperature:Pressure | -1.434 | 0.331 | -4.338 | <0.001 |
| Pooled *Myotis* | (Intercept) | -5.031 | 0.272 | -18.504 | <0.001 |
| Pooled *Myotis* | Temperature | 0.663 | 0.307 | 2.162 | 0.031 |
| Pooled *Myotis* | Pressure | 0.245 | 0.293 | 0.834 | 0.404 |
| Pooled *Myotis* | Temperature:Pressure | 0.726 | 0.337 | 2.156 | 0.031 |

**Fig 3.**
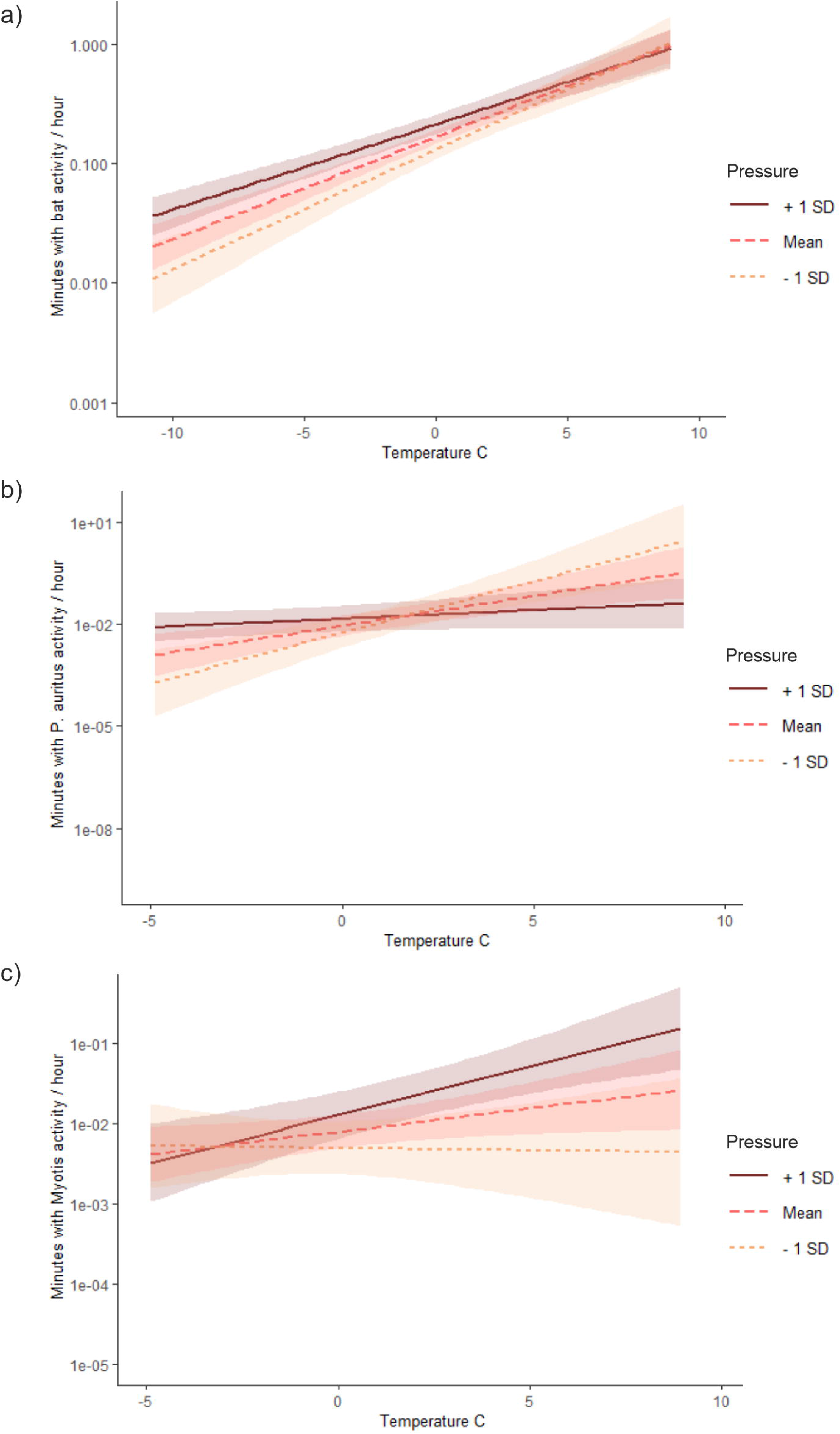
Effect of ambient temperature on a) the total winter bat activity (N=318 minutes with activity) b) activity of *Plecotus auritus* (N=66 minutes with activity) and c) *Myotis* species (Pooled for four species; *M. daubentonii*, *M. brandtii*, *M*. *mystacinus*, *M. nattereri*, N=35 minutes with activity) in different mean barometric pressures. Unit for activity is the predicted number of minutes with recordings / night. Shaded areas represent 95% confident intervals. Temperature and barometric pressure are mean values of previous 24 hours.

## DISCUSSION

Our results revealed greater bat activity at rock outcrops compared to root cellars, which have been considered one of the most important hibernacula in many areas of boreal zone (Rydell 1989; Lesiński et al. 2004; Vintulis and Pētersons 2014). Rock outcrops and boulder fields constitute up to 2.5 % of Finland’s land area (Kontula et al. 2018). Most of these are located in South-Western Finland, where our study was conducted, making them highly available as hibernacula. Almost all of the hibernation sites known in Finland prior to this study are manmade, but the number of individuals observed at these sites each winter likely resembles only a fraction of the true populations (Wermundsen and Siivonen 2010). Our results suggest that natural formations may be more important to hibernating bats than previously considered.

*Eptesicus nilssonii* were observed significantly more often than pooled *Myotis* species or *P. auritus*. The species is well adapted to the cold and long winters of the north, hibernating in colder environments than other species, leading to longer bouts of torpor that help them save more energy (Anufriev and Revin 2006; Siivonen and Wermundsen 2008). Furthermore, *E. nilssonii* have a higher body mass than *P. auritus* and *Myotis* species, which enables them with greater fat reserves for the winter (Rydell 1993). Larger body size further increases energy saving through relatively lower levels of heat loss, as their surface area to volume ratio is lower (Worthy and Edwards 1990). These factors may allow *E. nilssonii* to spend more energy on locomotion during the winter and permit them to forage when conditions are suitable. Lausen and Barclay (Lausen and Barclay 2006) reported a similar pattern of higher activity of *Eptesicus fuscus* compared to *Myotis* species during the winter in Canada.

While our results can indicate that *E. nilssonii* are more active during the winter compared to the other species monitored, it is important to note that this difference can be explained by other factors. Currently there are no estimates on population sizes of bats in Finland, and we therefore cannot determine whether our results are indeed reflecting differences in the activity or population size, or differences in hibernation habitat preferences. Furthermore, the acoustic detectability varies between the species. Detectability is influenced by the intensity of calls as well as the frequency, as high frequency sounds attenuate more quickly in the atmosphere (Griffin and Galambos 1941; Lawrence and Simmons 1982), leading to species such as *E. nilssonii* with high intensity and low frequency calls being more frequent in acoustic data compared to species with lower detectability (Schnitzler and Kalko 2001). Bat species also differ in their preference for abiotic conditions of the hibernacula and it is known that both *E. nilssonii* and *P. auritus* often hibernate in conditions that are colder and less humid than conditions preferred by many *Myotis* species (Wermundsen and Siivonen 2010). Given the ability of *E. nilssonii* to take advantage of hibernacula not suitable for other species (Masing and Lutsar 2007), it is also possible that the hibernacula studied here are not suitable for species requiring higher ambient humidity and temperature. We cannot disclose the possibility that other hibernation habitats, not included in our study, may be more suitable for *Myotis* species.

We predicted that ambient temperature would have a positive effect on bat activity, and that activity would be greater at low barometric pressure. In accordance with our hypothesis, we found that an increase in ambient temperature had a positive effect on the activity of *E. nilssonii*, *P. auritus* and *Myotis* species during the winter months. Furthermore, we found that total bat activity as well as the activity of *P. auritus* increased more rapidly with rising ambient temperature when barometric pressure was low. In Finland, low pressure systems during the winter often conjure mild temperatures together with cloudy, rainy weather, while high pressure systems usually arrive from the east, bringing cold yet clear weather (Similä 1981). We suggest that this result indicates that in general bats may indeed use low barometric pressure as a proxy for suitable conditions to leave the hibernaculum, leading them to be more active during warm, often cloudy, nights. Due to the siting of our passive detectors and the structure of most of the habitats studied, with no single clear entrance to the hibernacula, we were likely to only record bats that left the hibernaculum entirely and continued flying outside for a period of time. Because *P. auritus* is known to forage during the winter (Hays et al. 1992), we suggest that this species may benefit from detecting changes in barometric pressure from within the hibernaculum. In contrast, we found that the activity of *Myotis* species increased more rapidly with temperature in high barometric pressure, while the activity of *E. nilssonii* increased in high barometric pressure. We propose that these results show that the activity of *Myotis* species and *E nilssonii* is rather determined by the timing of their arousals than by foraging possibilities.

We suggest that a proportion of the winter activity we recorded was due to bats relocating to a different hibernaculum. Bats can benefit from timing these movements to warm nights, as this leads to reduced energy loss, corresponding with our model predicting more activity at warm ambient temperature (Klüg-Baerwald et al. 2016). We suggest that *E. nilssonii* may be more prone to shifting between hibernation sites due to its larger body size and fat reserves, as this enables more energy to be allocated for activity. Masing and Lutsar (2007) found that in case of severe frost reaching inside the hibernaculum, *E. nilssonii* were able to arouse and switch to another hibernaculum, whereas bats belonging to the genus *Myotis* generally froze to death. This is consistent with our result of *E. nilssonii* being active in colder temperatures than *Myotis* species.

In addition to their abundance, sites such as rock outcrop formations may offer additional advantages. For instance, it is likely that human disturbance is scarce, if nonexistent, in these environments compared to root cellars, bunkers, and other man-made structures. Furthermore, Michaelsen et al. (2013) suggested that using hibernacula such as rock scree might help bats avoid predators during hibernation and emergence. Despite the acoustic evidence of bat activity in many locations, we only found one hibernating bat in the natural study sites (*M. brandtii* inside a cave at a rock outcrop), which highlights the difficulty of visually observing hibernating bats in areas that lack large caves. Our results indicate that hibernation sites that are not accessible to humans, and have therefore been understudied, are more important for bats than has previously been acknowledged. Most research on hibernation behavior thus far has been conducted on species that hibernate in caves in karst regions, often in large numbers, leaving the majority of species not encountered in these environments understudied. Recently, the use of acoustic monitoring and studies utilizing temperature sensing radio tags has brought new insight to the hibernation behaviour of more elusive species (e.g. Lausen and Barclay 2006; Lemen et al. 2017; Ossa et al. 2020), to which our results add. However, climate change may affect the usability of these sites for bats, as the loss of snow cover is likely to have a great impact on the microclimates of these hibernacula. Furthermore, our results also highlight the need of taking bats into account in land use, when sites with rock outcrops are altered.

## ACKNOWLEDGEMENTS

The study was funded by Kone foundation, Emil Aaltonen foundation and H2020 Marie Sklodowska-Curie Actions. We thank Rauno Varjonen for maintaining the SongMeter in Vehmaa and Sari Haartonen from the Finnish Meteorological institute for her kind help in gathering the weather data.

